# Novel role of the lncRNA EPR as oncosuppressor in intestinal cancer

**DOI:** 10.64898/2026.04.28.719975

**Authors:** N Shim, M Rossi, M Nicolau, JR Barajas, E Zapparoli, P Briata, PL Puri, R Gherzi, L Caputo

**Author notes:** Authors for correspondence: Roberto Gherzi and Luca Caputo. These Authors contributed equally (N Shim, M Rossi).

## Abstract

We previously reported that the murine lncRNA *Epr* is essential for maintaining colon mucosal integrity and permeability. Mice lacking *Epr* in the colon are more susceptible to colitis and tumor development. Additionally, we demonstrated that human *EPR* expression is reduced in ulcerative colitis and in a small cohort of colon adenocarcinoma patients. Here, we present evidence that human and mouse *EPR* share several key physiological features: preferential binding to the KH1 domain of their interacting protein, KSRP; specific expression in canonical and immature goblet cells of the large intestine; and a functional role in intestinal goblet cell development. The correlation between *EPR* levels and survival in large cohorts of metastatic colon adenocarcinoma patients, together with the capacity of human *EPR* to inhibit cell proliferation and induce apoptosis in two distinct human colon adenocarcinoma cell lines, suggests that *EPR* may serve as both a valuable prognostic marker for goblet cell-derived adenocarcinomas and a potential therapeutic target.

## INTRODUCTION

Long non-coding RNAs (lncRNAs) regulate gene expression at multiple levels by influencing chromatin remodeling and coordinating transcriptional and post-transcriptional processes through direct interactions with nucleic acids or by associating with DNA- or RNA-binding proteins (Mattick et al. 2023). LncRNA expression is typically tissue-specific and developmentally regulated, with even a small number of copies having significant regulatory effects (Nemeth et al. 2024; Mattick et al. 2023). Most lncRNAs are species- or lineage-specific, with only a small fraction showing detectable sequence conservation across distant species (Mattick et al. 2023; Zhou et al. 2023). When present, conservation often centers on promoters, short exonic motifs, or RNA-binding protein (RBP) sites (Washietl et al. 2014). In fact, for lncRNAs, conservation is best detected through a combination of structural and functional analyses, as many are overlooked by sequence alignment alone. Since the identification of the first lncRNA, *H19* (Pachnis et al. 1984), the number of lncRNAs identified and the roles of this class of regulatory RNA expanded, ranging from development to cancer (reviewed in (Perry and Ulitsky 2016; Peng et al. 2017; Huarte 2015; Delás and Hannon 2017). Due to the heterogeneity of lncRNAs, their role in cancer progression is transcript specific. While, for example higher expression of lncRNAs *HOTAIR* and *ANRIL* in tumors is associated to metastatic stage and poor prognosis (Gupta et al. 2010; Kotake et al. 2011), the lncRNAs *NBAT1, PR-lncRNA-1* or *PTENP1* inhibit cells proliferation and their expression is associated with better clinical outcomes (Pandey et al. 2014; Sánchez et al. 2014; Poliseno et al. 2010). Recently, the role of lncRNAs in maintaining intestinal health has been extensively studied (Xiao et al. 2019; Lu et al. 2023).

We recently reported that the lncRNA *Epr* (BC030870, ENSMUSG00000074300) regulates mucus production, intestinal permeability, and susceptibility to inflammatory stimuli, thereby supporting the intestinal chemical barrier by modulating the transcription of specific target genes in mice (Briata et al. 2023). Furthermore, our findings suggest that *Epr* is involved in colon cancer development in mice and in human colon carcinogenesis (Briata et al. 2023). The human equivalent of mouse *Epr* (LINC01207, also known as SMIM31 or ENSG00000248771) has not been extensively studied in the context of intestinal physiology and pathology. We observed a decrease in human *EPR* expression in datasets derived from RNA-Seq analyses of human colon biopsies from patients with ulcerative colitis, compared to healthy controls (Briata et al. 2023). A similar reduction was also observed in colon adenocarcinoma samples from a small patient cohort (Briata et al. 2023).

Although mouse and human *EPR* share some sequence similarities, neither primary sequence analysis nor in silico assessment of their potential secondary structures revealed clear similarities. This study aims to explore structural and functional conservation between mouse and human *EPR* by: (i) analyzing how the lncRNA interacts with KSRP, a confirmed mouse *Epr* binding protein (Rossi et al. 2019); (ii) investigating *EPR* expression in human colon mucosa; (iii) assessing *EPR* expression dynamics during organ development using human induced pluripotent stem cell (iPSC)- derived organoids; and (iv) examining *EPR*’s function in human colon cancer cells.

Our results demonstrate that: (i) both human and mouse *EPR* interact similarly with KSRP; (ii) both lncRNAs are primarily expressed in goblet cells of the small and large intestines; (iii) human *EPR* begins expression at the mid-hindgut stage and peaks in late organoids, roughly mirroring what is observed in mouse embryos; (iv) patient survival rates in colon cancer are positively correlated with *EPR* expression, and *EPR* levels tend to be higher in less advanced cases; and (v) ectopic expression of *EPR* in human colon cancer cells reduces their proliferation.

## RESULTS AND DISCUSSION

### Human and murine Epr exhibit comparable interactions with KSRP

Originally, we identified mouse *Epr* as a lncRNA that interacts with the multifunctional nucleic acid-binding protein KSRP (also known as FUBP2) in a genome-wide screening conducted by anti-KSRP ribonucleoprotein complex immunoprecipitation followed by RNA sequencing (RIP-Seq) in normal murine mammary gland cells (Rossi et al. 2019). This interaction was confirmed through band-shift analysis (Rossi et al. 2019). Further, KSRP and mouse EPR functionally interact to control cell proliferation (Rossi et al. 2019). Sequence alignment of mouse and human *EPR* shows an overall 70% identity in the region from nucleotide (nt) 3 to nt 535 of human *EPR* (**Supplementary Figure S1**), but does not show any similarity in its 3’ region, and the ability of human *EPR* to interact with KSRP has never been experimentally tested.

As a first step toward gaining insights into the role of human *EPR* in large intestine physiology and pathology, we investigated the potential interaction between human *EPR* and its putative binding protein, KSRP. To this end, we utilized the AlphaFold Server, a web-based tool powered by the AlphaFold 3 AI model, which generates 3D structure predictions for biomolecular complexes (Abramson et al. 2024). The analysis of full-length KSRP and human *EPR* complex prompted us to focus further analyses on the four KH domains of KSRP in complex with the 523 most 5’ nt of human *EPR* (data not shown). These results were predictable since it has been extensively reported that the four KH domains are the structures responsible for KSRP interaction with a variety of nucleic acids (Gherzi et al. 2014), and, on the other hand, the effects of full-length mouse EPR on cell proliferation in mammary gland cells were reproduced by a deletion mutant encompassing nt 1-760 of mouse *Epr* (*Briata and Gherzi, unpublished observation)*. This second analysis allowed us to further restrict our investigation to a region of human *EPR* encompassing nt 321-393 and a mouse *Epr* region from nt 321-386 (data not shown). Finally, we further restricted our analysis to nt 384-392 (human *EPR*) and nt 353-363 (mouse *EPR*), and modeled the interactions of these RNA sequences with each individual KH domain of either human or mouse KSRP (**Figure 1A**). As a metric of overall prediction accuracy, we focused on two parameters: pTM (predicted Template Modeling score) as an index of overall complex accuracy and ipTM (interface predicted Template Modeling score) as an estimate of the accuracy of the protein-RNA interaction interfaces within the structure. Both metrics are scalar values between 0 and 1, where higher values indicate a more confident and likely correct prediction (Abramson et al. 2024). pTM is particularly useful for evaluating the interaction accuracy of the entire predicted protein/RNA complexes. At the same time, ipTM assesses the confidence in how the different parts of the complex fit together at the interface. Both scores must be used together to obtain a reliable evaluation of the quality of the prediction, as a complex with a good pTM but low ipTM may have a correct global fold but a poor or incorrect interface.

**Figure 1.**
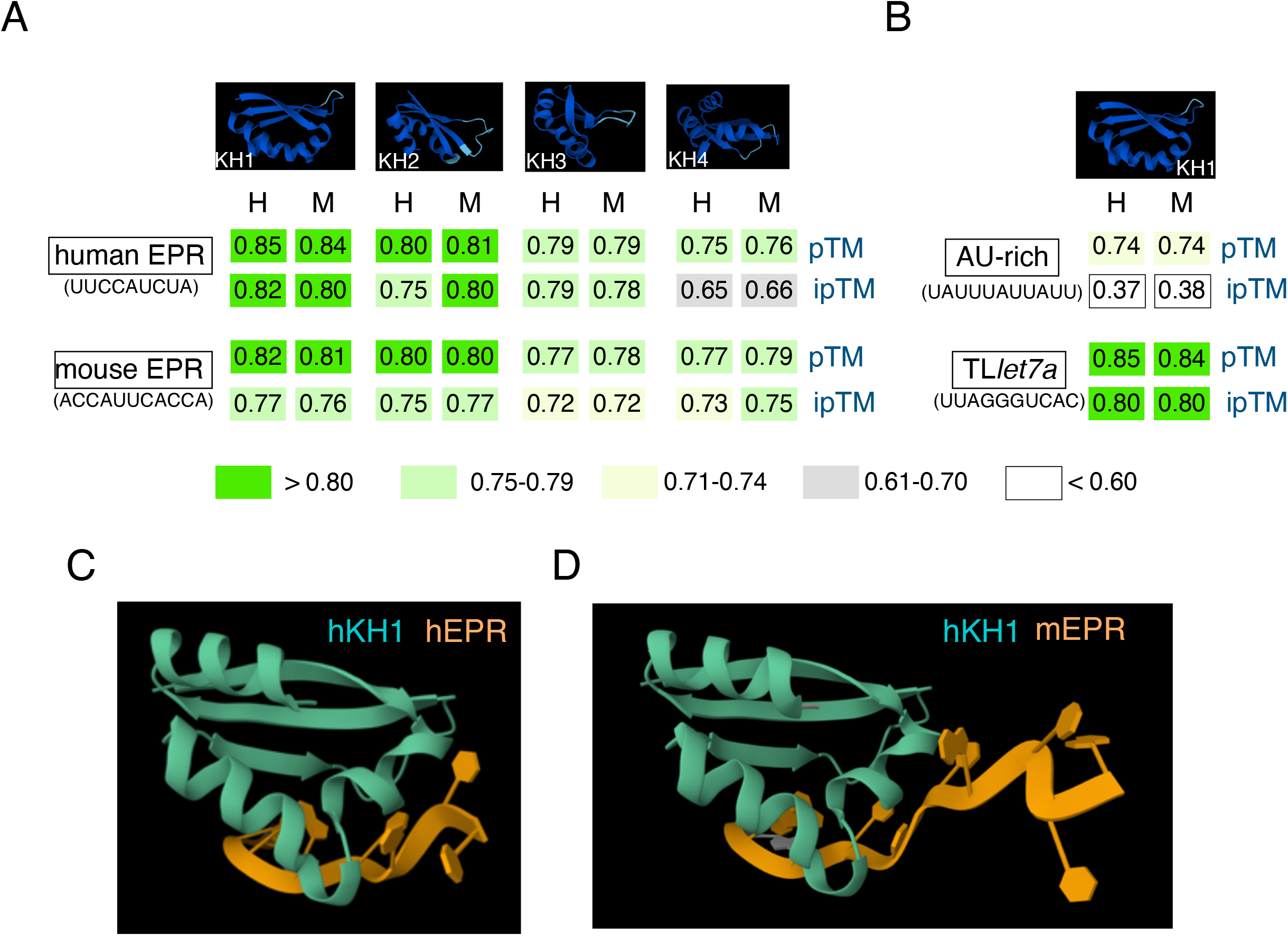
Human and murine EPR exhibit comparable interactions with KSRP. (A) Top, predicted 3D structure of each of the four KH domains that form the nucleic acid binding regions of human KSRP; bottom, pTM and iPTM values calculated for the interaction of each domain with either human (H) or murine (M) EPR. (B) Top, predicted 3D structure of the KH1 domain of human KSRP; bottom, pTM and iPTM values calculated for the interaction of each of the human (H) or murine (M) KH domains with the AU-rich motif (García-Mayoral et al., 2007) or the terminal loop of pri-let-7a (TLlet7a, Trabucchi et al., 2009). (C, D) predicted structures of the KH1 domain of human KSRP in association with human EPR and mouse Epr, respectively.

Our data indicate that either the human or mouse KH1 and KH2 domains of KSRP interact with human and mouse *EPR*, with KH1 exhibiting a stronger interaction capacity than KH2 (**Figure 1A**). Conversely, KH3 and KH4 interactions with either human or mouse *EPR* do not conform to the prediction criteria we established (**Figure 1A**). As positive and negative controls, we evaluated the interaction of KH1 with the terminal loop of pri-*Let7a* (TL*let7a*) (Trabucchi et al. 2009) and with an AU-rich sequence (García-Mayoral et al. 2007), respectively. As anticipated, KH1 demonstrates a robust interaction with TL*let7a*, while the modeling of KH1 with the AU-rich sequence fails to meet the criteria for a reliable interaction (**Figure 1B**). Finally, we predicted the structures of the KH1 domain of human KSRP in association with human and mouse *EPR*, respectively, showing that *EPR* interacts with the alpha-helix structure present in KH1 (**Figure 1C** and **1D**).

In summary, these data suggest that both human and mouse *EPR* exhibit comparable affinity for the KH1 domain of KSRP.

### *EPR* expression in human intestinal organoids

The expression of *Epr* is first detected at E7.5 in the mouse embryo (Rossi et al. 2019). At this developmental stage, the precursors of the large intestine are not yet morphologically distinct structures. Instead, they form part of the posterior endoderm, a layer of cells that develops during gastrulation (Wells and Spence 2014). At this stage, the definitive endoderm (DE) emerges from the posterior end of the primitive streak. DE is influenced by signaling interactions with the posterior mesoderm, a process that establishes the anterior-posterior body axis (Wells and Spence 2014). Interestingly, *Cdx2* is expressed in the hindgut and posterior primitive streak of the mouse embryo around 7.5 to 9.5 days postcoitum (Wells and Spence 2014). Analysis of ChIP-seq datasets from human colonic cell lines revealed that *CDX2* associates with the *EPR* promoter region (**Supplementary Figure S2A**) (Meyer et al. 2012).

To explore the role of *EPR* in the development of the human large intestine, we used human induced pluripotent stem cell (iPSC)-derived colonic organoids (Spence et al. 2011). By applying precise combinations of growth factors and signaling molecules, human iPSCs can be differentiated into definitive endoderm, hindgut, and, ultimately, colonic tissue, forming intestinal organoids that mimic large-intestine development. Single-cell RNA-sequencing (scRNA-seq) shows that iPSC-derived colonic organoids closely resemble the fetal human colon, recapitulating key gene expression programs and regional identities (Xu et al. 2025).

Human iPSCs underwent a well-defined differentiation protocol, with cultures analyzed at the indicated time points (**Figure 2A**). Representative brightfield and immunofluorescence images of hiPSCs undergoing differentiation into intestinal organoids are shown in **Figure 2B** and **Supplementary Figure S2B**. We examined the expression of several genes that mark different stages of organoid differentiation. As expected, the expression of *OCT4*, a factor that maintains pluripotency, is rapidly downregulated during differentiation, while *SOX17*, which encodes a key regulator of definitive endoderm specification and indicates successful exit from pluripotency and commitment to endodermal fate, is expressed after definitive endoderm formation (**Figure 2C**). The expression of *CDX2*, which encodes a master transcription factor for intestinal identity necessary for specifying and maintaining intestinal fate, and *LGR5*, a marker of intestinal stem cells essential for crypt formation, self-renewal, and multilineage differentiation, mirrors the expression of *EPR* and its target gene *MUC2* (**Figure 2D**).

**Figure 2.**
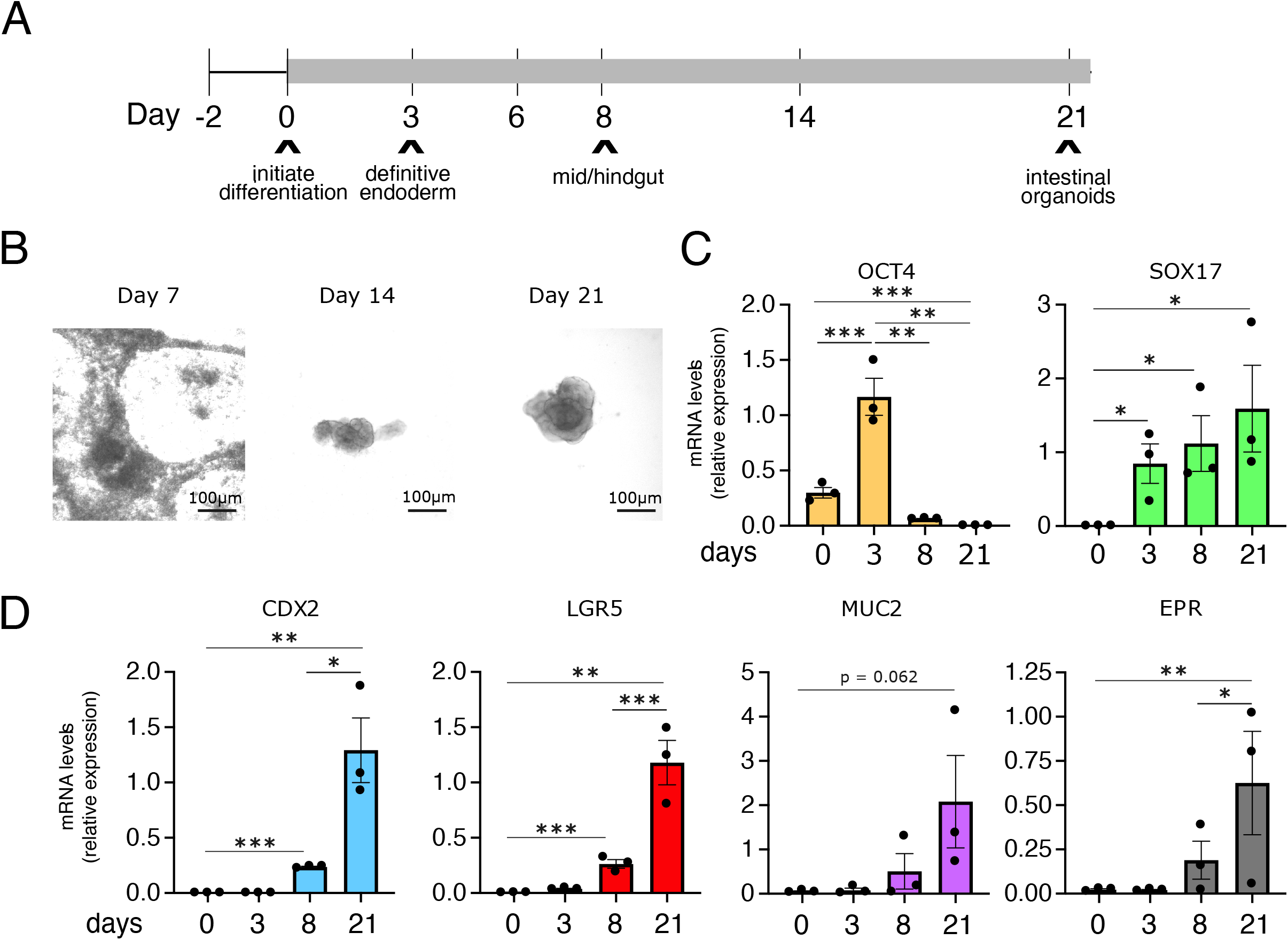
EPR expression in human intestinal organoids. (A) Timeline of the stage of differentiation of human iPSCs upon culture with specialized media to generate human intestinal organoids (see Materials and Methods). (B) Representative brightfield images of intestinal organoids cultures derived from hiPSCs at different stages (days) of the differentiation protocol. Scale bar 100µm. (C, D) qRT– PCR analysis of stage-specific pluripotent, endodermal and intestinal markers at multiple stages (days) of the differentiation protocol. Expression levels have been normalized over *ACTB* expression. The values are averages (±SEM) of three independent experiments performed in triplicate. * p≤0.05, ** p≤0.01, ***p≤0.001

Overall, these data suggest that *EPR* is a target of CDX2 and that it plays a role in the specification of intestinal goblet cells.

### Human *EPR* marks canonical goblet cells in the large intestine

*EPR* is mainly found in the small and large intestines of humans (**Supplementary Figure S3A**), as observed in mice (Rossi et al. 2019; Briata et al. 2023). scRNA-seq has shown that goblet cells (GCs) have the highest levels of *EPR* expression (**Figure 3A**), consistent with our findings in mice (Wang et al. 2019; Briata et al. 2023).

**Figure 3.**
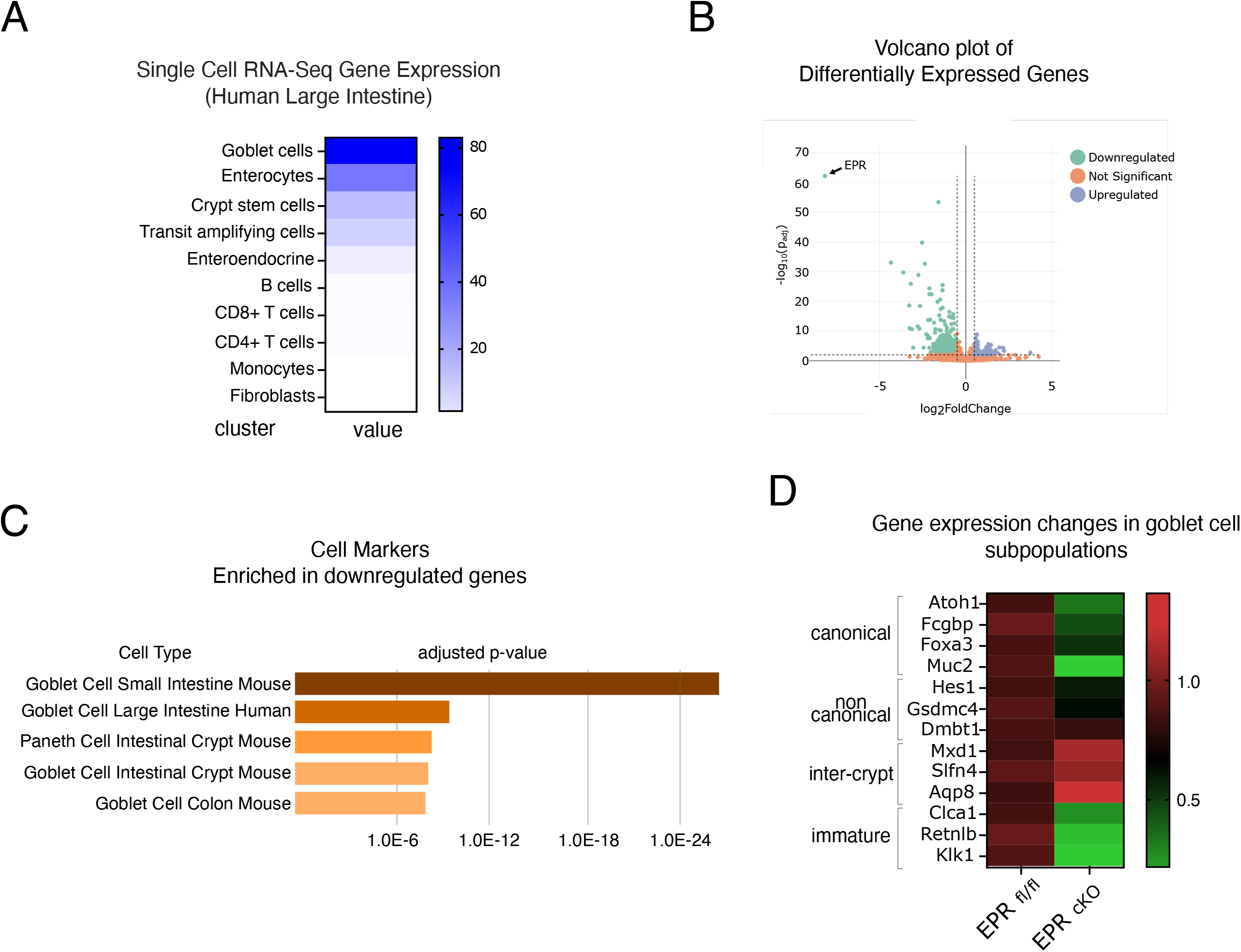
Human EPR marks canonical goblet cells in the large intestine. (A) Heat map of the single-cell RNA-Seq data from the human large intestine, retrieved from Wang et al., 2020. (B) Volcano plot showing differentially expressed genes in colon crypts from either *Epr*^*fl/fl*^ or *Epr*^*cKO*^ mice, re-analyzed with a cutoff of |log2FC| > 0.5. (C) Gene Ontology cell marker analysis (using the online EnrichR tool) of transcripts that are downregulated in *Epr*^*cKO*^. (D) Heat map illustrating changes in the expression of genes specific to each goblet cell subpopulation in *Epr*^*cKO*^ versus *Epr*^*fl/fl*^ RNA sequencing.

We previously demonstrated that expression of *Muc2, Fcgbp*, and *Claca1* genes, which encode the core component of mucus produced by GCs, is significantly reduced in mice in which *Epr* was deleted using a CRE recombinase under the control of *Cdx2* promoter, a transcription factor instrumental in the development and maintenance of the intestinal epithelium, leading to specific deletion of *Epr* in the epithelium of the distal ileum, cecum and throughout the colon from the crypt base to the luminal surface (*Epr*^*cKO*^) (Briata et al. 2023). We were intrigued by evidence that, despite this reduction, GCs are still present in the large intestine of *Epr*^*cKO*^ mice (Briata et al. 2023). Here, we reanalyzed our RNA-Seq data using a cutoff of |log_2_FoldChange (FC)| > 0.5. The new analysis confirmed that the majority of genes are down-regulated in *Epr*^*cKO*^ (**Figure 3B**), and that GCs are the main cell population affected by *Epr* deletion in the colon (**Figure 3C**). Over the past decade, GCs have emerged as key players in establishing the protective mucus barrier that covers the intestine and in modulating gut immune responses by sampling and delivering luminal antigens to the immune system, thereby aiding the induction of adaptive immune responses (Birchenough et al. 2015). scRNA-seq trajectory analysis of *Muc2*-expressing cells in mouse and human revealed that GCs separate into distinct trajectories marked by specific markers: canonical GCs, non-canonical GCs, and inter-crypt GCs (Nyström et al. 2021; Hickey et al. 2023; Burclaff 2023). Our data show that *Epr* gene deletion in mice significantly reduces canonical GCs, as evidenced by a marked decrease in their markers *Atoh1, Fcgbp, Foxa3* and *Muc2* (**Figure 3D**). However, *Epr*^*cKO*^ does not impact non-canonical GCs, with no significant changes observed in *Hes1, Gsdmc4, Dmbt1*, or inter-crypt GCs, with no notable differences in *Mxd1, Sfln4*, and *Aqp8* (**Figure 3D**). Additionally, *Clca1, Klk1*, and *Retnlb*, markers for human immature goblet cells (Hickey et al. 2023), are significantly reduced in the absence of *Epr* (**Figure 3D**). Our reevaluation of RNA-seq data also revealed a notable downregulation of the *Best2* gene in *Epr*^*cKO*^ mice (log_2_FC = -1.34, p = 0.0048). *Best2* is a marker for a subset of colorectal-specific mature goblet cells critical for bicarbonate-mediated mucus secretion (Yu et al. 2010). *BEST2*-positive goblet cells are specifically decreased in active ulcerative colitis lesions (Ito et al. 2013), and lower *BEST2* expression correlates with shorter survival in colon cancer patients (Wang et al. 2022). Lastly, the data indicating that *Sox9, Rgmb, Smoc2, Lgr5*, and *Ascl2* are not significantly affected in *Epr*^*cKO*^ mice (**Supplementary Figure S3B**) rules out the possibility that *Epr* influences the colon stem cell lineage.

Our past and present data support a role for *EPR* in the function of immature and canonical subpopulations of colonic goblet cells, leading us to hypothesize that *EPR* reduction or dysfunction may affect these cells in humans, resulting in pathological conditions, including Inflammatory Bowel Disease and colon cancer.

### Colon cancer patients with higher *EPR* expression have a longer survival

The literature reports conflicting findings regarding *EPR* expression in colorectal cancers. Our study revealed a significant reduction in *EPR* expression in cancer tissue compared with adjacent normal tissue in patients with colon adenocarcinoma (Briata et al. 2023). In this report, we refined our analysis and now demonstrate that *EPR* expression is significantly higher in Stage I and II adenocarcinoma compared to Stage III and IV (**Figure 4A**). Subsequently, we extracted survival data from a large cohort of colon cancer patients in the Genomic Data Commons Data Portal, maintained by the National Cancer Institute. Based on TPM (transcripts per million) values for each gene, patients were categorized into two groups according to their expression levels (cut-off value: 0.22), and the correlation between expression and patient survival was examined. The prognosis of each group was assessed using Kaplan-Meier survival estimators, and the survival outcomes of the two groups were compared using log-rank tests. Patients with higher *EPR* levels have a longer survival probability than those with lower *EPR* levels. Furthermore, the statistical significance of the survival difference is greater in Stage III and IV patients (**Figure 4B**).

**Figure 4.**
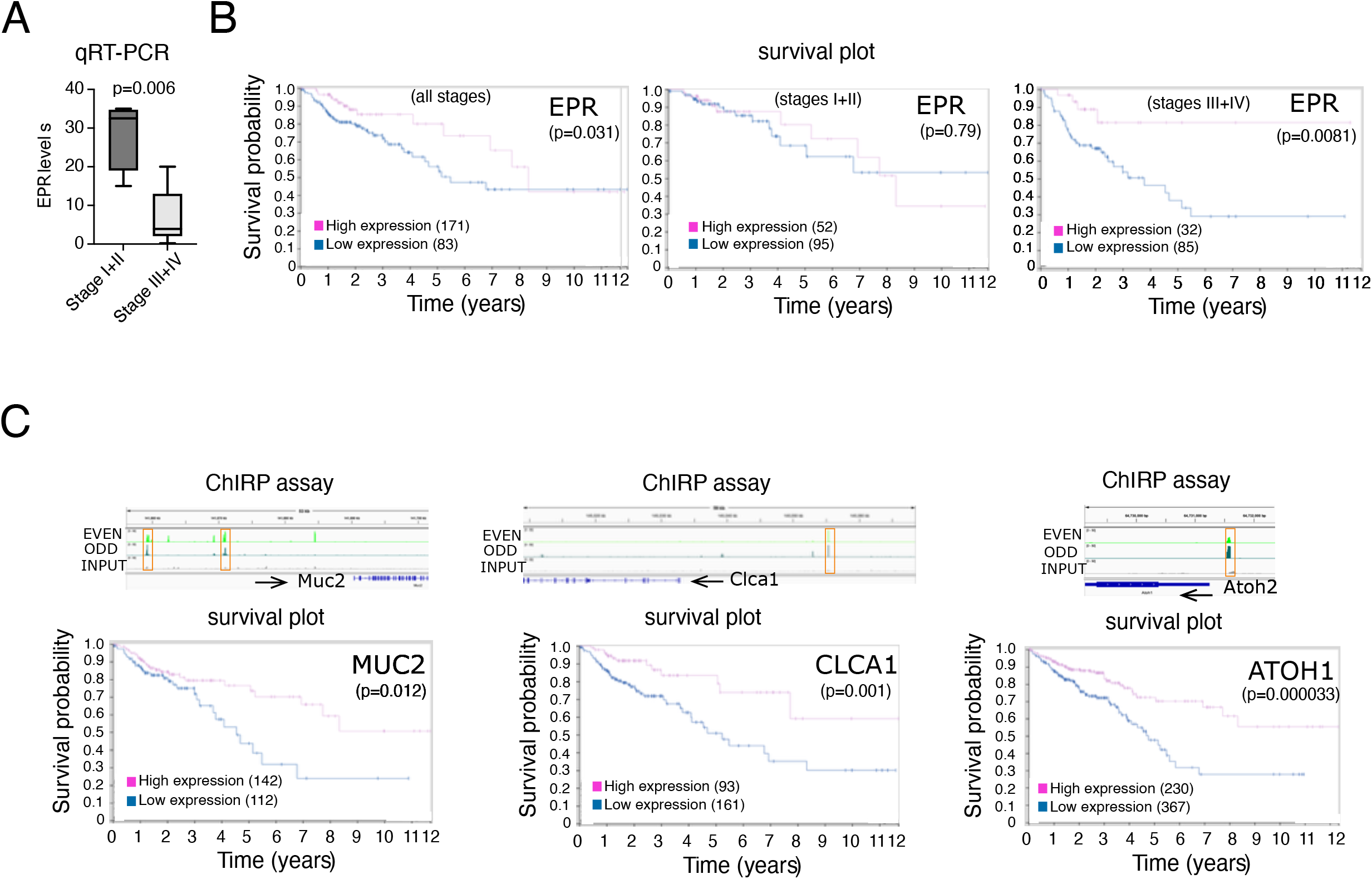
Patients with colon cancer who have higher EPR expression tend to have a longer survival rate over time. (A) The expression of *EPR* was analyzed by qRT-PCR in the cohort of colon adenocarcinomas and adjacent normal tissue described in Briata et al. (2023). Tumors were grouped according to their stage, and the statistical significance (Student’s t-test) of the expression differences between Stages I+II and Stages III+IV is presented. (B) Survival analysis in large cohorts of patients with colon adenocarcinoma grouped according to their clinical stages. (C) Top panels, association of *Epr* with the chromatin at the indicated genomic loci as assessed by ChIRP analysis (Zapparoli et al., 2020). Bottom panels, survival analysis in patients with colon adenocarcinoma. P is the log-rank value for the Kaplan-Meier plots showing results from the analysis of correlations between the expression levels of *EPR* (B) and its target genes (as indicated in C) and patient survival.

To gain further insight into the importance of *EPR* expression in colon cancer prognosis, we examined survival outcomes based on the expression levels of *EPR*’s direct target genes that we previously identified (Briata et al. 2023). ChIRP (Chromatin Isolation by RNA Purification) analysis conducted in mouse colon cells identified direct binding of *EPR* to potential regulatory elements in the promoter region (5’ to the transcription start site) of *MUC2, CLCA1*, and *ATOH1* (**Figure 4C**, top panels). Higher levels of *MUC2, CLCA1*, and *ATOH1* are associated with longer survival (**Figure 4C**, bottom panels). Our analyses indicate that EPR downregulation is more pronounced in severe colon cancer cases and is associated with a worse prognosis.

### *EPR* overexpression impairs proliferation and increases CDKN1A expression in colon cancer cells

Previously, we described mouse *Epr* as a lncRNA that inhibits cell proliferation and increases the expression of the cyclin-dependent kinase inhibitor CDKN1A (also known as p21WAF1/Cip1) in both immortalized and transformed mammary cell lines, as well as in an animal model of orthotopic tumor transplantation (Rossi et al. 2019). Additionally, cell cycle analysis of immortalized mammary gland cells overexpressing *EPR* showed a significant increase in cells arrested in the G1 phase (Rossi et al. 2019). When we ectopically expressed *EPR* in the colon cancer cell line SW480, we observed higher levels of cleaved Caspase-3 and increased expression of pro-apoptotic genes, although we did not observe any changes in CDKN1A expression or cell cycle progression (Inga et al., *unpublished observation*). Interestingly, SW480 cells do not produce detectable levels of the *EPR* gene (Briata et al. 2023). In this report, we examined *EPR* function in transfected colon cancer cell lines that either do or do not express endogenous *EPR*.

We performed a preliminary cell screening based on the co-occurrence of H3K4me3 and H3K27ac marks at the promoter of the *EPR* gene across different colon cancer cell lines (Zheng et al. 2019). The co-occurrence of H3K4me3 and H3K27ac is a hallmark of active promoters (Beacon et al. 2021). We selected HCT-8 and LS180 as cells in which the *EPR* promoter is active, and HCT-116 as an *EPR* promoter-inactive cell line, respectively (**Supplementary Figure S4A**). The expression of *EPR* in HCT-8 and LS180, but not in HCT-116 was confirmed by qRT-PCR (**Supplementary Figure S4B**).

We transfected either HTC-8 or HCT-116 cells with the empty pBICEP-CMV-2 vector (HCT-8 mock, HCT-116 mock) or with human *EPR* cloned into the pBICEP-CMV-2 vector (HCT-8 EPR, HCT-116 EPR), then analyzed the transfectants 48 hours post-transfection. *EPR* overexpression in transfected cells was confirmed by qRT-PCR (**Supplementary Figure S4C**). First, we evaluated the effect of EPR overexpression on HCT-8 cell proliferation by measuring the expression of the proliferation marker Ki-67. Both *MKI67* transcript and Ki-67 protein are consistently upregulated in colon adenocarcinoma tissues compared to adjacent normal colon tissues (Clinical Proteomic Tumor Analysis Consortium). EPR overexpression significantly reduced Ki-67 expression in HCT-8 cells but had no effect in HCT-116 cells (**Figure 5A** and data not shown). Previously, we observed that overexpression of mouse *Epr* in SW480 cells induces the expression of positive regulators of programmed cell death (Briata et al. 2023). Additionally, we found that cleaved Caspase-3 levels are increased in cells expressing mouse EPR (Briata et al. 2023). Importantly, the expression of CDKN1A is increased in HCT-8 cells 48 hours after transfection with human *EPR* (**Figure 5B**).

**Figure 5.**
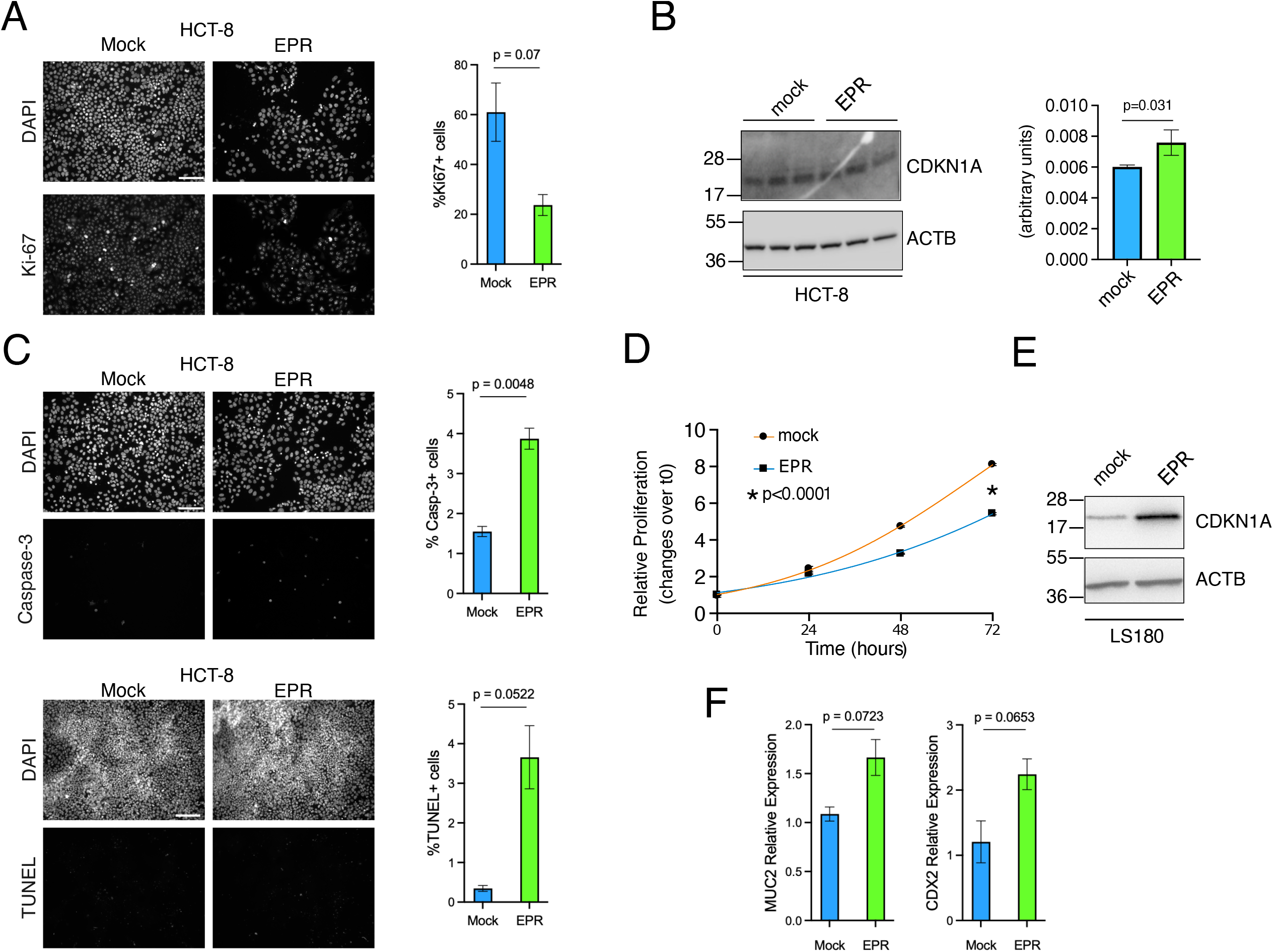
EPR overexpression impairs proliferation and enhances CDKN1A expression in colon cancer cells. (A) Immunofluorescence analysis of Ki-67 in HCT-8 cells transfected with empty plasmid (mock) or EPR expressing plasmid (EPR) and quantification (right). Scale bar 200µm. (B) CDKN1A protein levels in HCT-8 transfected with empty plasmid (mock) or EPR expressing plasmid (EPR). (C) Immunofluorescence analysis of cleaved Caspase-3 (top panels) and TUNEL assay (Bottom panels) in HCT-8 cells transfected with empty plasmid (mock) or EPR expressing plasmid (EPR) and quantification (right). Scale bar 200µm. (D) Cell proliferation analysis (using Crystal violet staining) of either mock or EPR (EPR) -overexpressing LS180 cells. (E) Immunoblot analysis of total cell extracts from either mock or EPR-overexpressing LS180 cells. (F) qRT-PCR expression analysis of MUC2 and CDX2 in HCT-8 cells transfected with empty plasmid (mock) or EPR expressing plasmid (EPR). The values are averages (±SEM) of three independent experiments performed in triplicate.

We analyzed the activation of the apoptotic pathway in human *EPR*-transfected HCT-8 cells compared with mock-transfected cells. We observed increased positive staining for both anti-cleaved Caspase-3 and TUNEL assays upon EPR transfection (**Figure 5C**).

To confirm that the EPR-mediated induction of apoptotic program is not restricted to HCT-8 cells (a male cell line, negative for KRAS^G12D^ mutation), we transfected an unrelated colon carcinoma cell line, LS180 cells (a female cell line, positive for KRAS^G12D^ mutation), which express a moderate amount of endogenous *EPR*, with either human *EPR* or an empty vector. The overexpression of *EPR* in transfected LS180 cells was confirmed by qRT-PCR (**Supplementary Figure S4D**). Human *EPR* significantly reduced LS180 cell proliferation (**Figure 5D**) and increased the expression of the cyclin-dependent kinase inhibitor CDKN1A (**Figure 5E**). Interestingly, the expression of *MUC2*, a prominent target of *EPR* in mice (Briata et al. 2023) (see also **Figure 4C**), and of the differentiation marker CDX2 are elevated by transfecting human *EPR* into HCT-8 cells (**Figure 5F**), suggesting a push towards more differentiated goblet-like cells.

In summary, similarly to murine *Epr* in colon cell lines (Briata et al. 2023) and in mammary gland cells (Rossi et al. 2019), human *EPR* inhibits cell proliferation, increases the expression of the negative regulator of cell cycle progression, CDKN1A, in colon cancer cells, and activates the apoptotic pathway.

## CONCLUSION

In sum, our study expanded on the biology of the human lncRNA *EPR* (LINC01207, also known as SMIM31 or ENSG00000248771) in the context of intestinal physiology and pathology. Using an organoid system, we confirmed *EPR* expression during early colon development at a stage compatible with the appearance of GCs and colon crypt specification. This finding was further supported by our murine model and scRNA-seq expression analysis, linking *EPR* to GC specification. Furthermore, building on previous findings in mammary gland tissues and murine models, we suggest that *EPR* could function as a novel oncosuppressor in human colon carcinoma. Analysis of publicly available datasets demonstrated a correlation between high *EPR* expression and longer survival among stage III+IV patients with colon adenocarcinoma. Finally, overexpression of *EPR* in two separate colon adenocarcinoma cell lines (HCT-8, negative for KRAS^G12D^, and LS180, positive for KRAS^G12D^) indicated that *EPR* induces a cell-cycle block, apoptosis, and ultimately, differentiation toward GCs. During the final preparation of this preprint, Wang and colleagues reported on the role of the lncRNA *AC007637*.*1* in regulating the Trim25/PCNA axis and inhibiting tumorigenesis in colorectal cancer (Wang et al. 2026). With the advancement of *in vivo* mRNA delivery using lipoparticles, it is tempting to speculate on a potential clinical use of *EPR* to treat late-stage colon adenocarcinomas.

## ACKNOWLEDGEMENTS

Work in Dr. Gherzi’s laboratory was supported by Associazione Italiana per la Ricerca sul Cancro (AIRC; I.G. 21541). Work in Dr. Puri’s laboratory was supported by NIAMS (R01 AR056712) and NIGMS (R01 GM13471). NS is supported by CIRM SPARK fellowship (EDUC3-13123). MN is supported by CIRM pre-doctoral fellowships (EDUC4-12813). We would like to thank all members of the Puri and Sacco laboratories for scientific discussion.

## AUTHORS’ CONTRIBUTIONS

LC and RG conceived the study, designed the experiments and wrote the manuscript. NS, MR, JRB, MN, EZ, PB performed the experiments. NS analyzed the RNAseq dataset with help from MN. PLP and RG provided funding support. All authors participated in the scientific discussion and in the review and editing of the manuscript. The authors read and approved the final manuscript.

## MATERIALS AND METHODS

### Cell Culture maintenance and culture

Colon cancer cell lines were grown in DMEM high glucose (Gibco Cat #1195092) supplemented with 10% FBS (Cell Culture Collective Cat #FB-11), and 1x Pen/Strep (Gibco Cat #10378016). Cells were passaged 1:5 when 80% confluent.

### Plasmid Transfection

Colon cancer cell lines were grown in complete media until approximately 50-60% confluent and transfected with EPR expressing plasmid, or empty control plasmid using Lipofectamine 3000 (Life Technologies L3000008) according to the manufacturer’s instructions. Cells were collected for downstream analysis 48 hours after transfection.

### Intestinal Organoids

Human embryonic stem cells (H9 - WiCell WA09) were maintained in mTESR+ media (Stem Cell Technologies Cat #100-0276) on Matrigel® (Corning Cat #354277) coated dishes in incubator at 37°C, 5% CO_2_. H9 cells were differentiated to small intestinal organoids using the STEMdiff Intestinal organoids Kit (Stem Cell Technologies Cat #05140) according to the manufacturer’s protocol.

### Quantification of cell proliferation by crystal violet

For some experiments, cell proliferation was assessed by crystal violet staining. At the indicated time after plating, cells were fixed (10% formalin) and stained with crystal violet solution (0.1% crystal violet in acetic acid). After two washes with water, crystal violet staining was measured by a spectrophotometer at a wavelength of 590 nm.

### RNA-seq analysis

Previously generated RNA sequencing (RNA-seq) dataset (Briata et al. 2023) has been further analyzed as follows. Raw counts have been downloaded from GEO with accession number GSE218994. Differential expression analysis was carried out with GENAVi (Reyes et al. 2019), using DESeq2 (Love et al. 2014). Differentially expressed genes in *Epr*^*cKO*^ versus *Epr*^*fl/fl*^ were filtered for |log2FC| > 0.5 and Benjamini-Hochberg corrected p-value (p_adj_) < 0.01. Enrichment analysis and gene ontology were performed using the online software EnrichR (https://maayanlab.cloud/Enrichr).

### Gene expression analysis

Total RNA was isolated using Quick-RNA™ Microprep Kit (Zymo Research Cat #R1050) according to manufacturer’s recommendation. 0.5-1 µg of total RNA was retrotranscribed using High-Capacity cDNA Reverse Transcription Kit (Applied Biosystems Cat #4368813) and qRT-PCR was performed using the Power SYBR Green PCR Master Mix (Applied Biosystems Cat #4367659) on Applied Biosystems 7900HT Fast Real-Time PCR. Relative gene expression was calculated using the 2^-ΔΔCT^ method and normalized to *TBP* or *ß-actin* expression, as defined in the figure legend. Primer pairs are shown below:

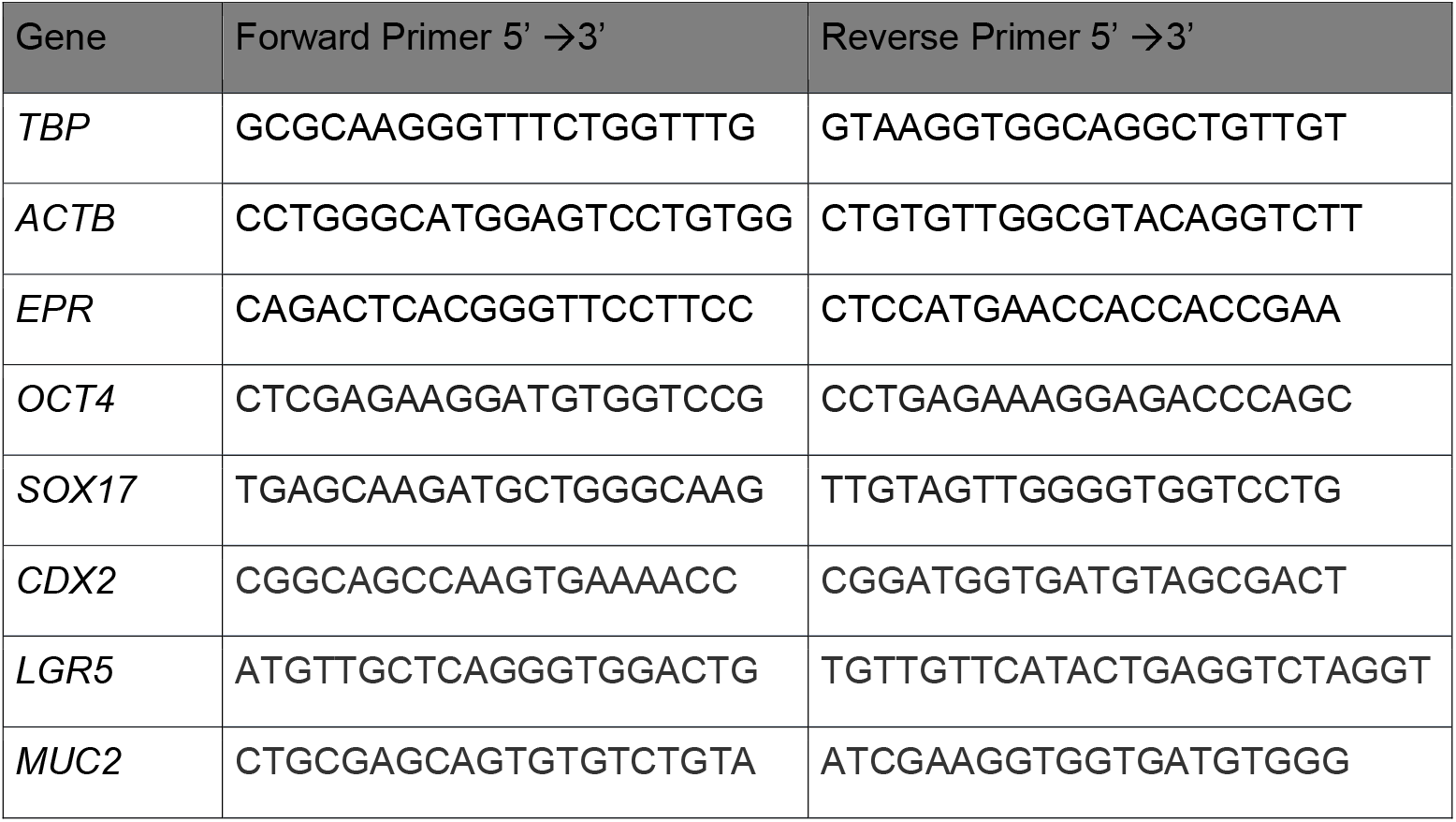

### Immunofluorescence staining

HCT-8 cells grown on 96-well tissue culture plates (Greiner Bio-One) were fixed with 4% PFA for 10 minutes at room temperature and permeabilized in 0.5% Triton X-100 for 10 minutes, then blocked in blocking buffer (4% BSA, 0.5% Triton X-100 in PBS) for 30 minutes. Incubation with the primary antibodies diluted in blocking buffer was performed overnight. Primary antibodies: rabbit anti-CDX2 (Cell Signaling Technology Cat #12306T), rabbit anti-Ki-67 (Abcam ab15580), rabbit anti-Cleaved Caspase-3 (Cell Signaling Technology Cat #9661S). Secondary antibodies: AlexaFluor goat anti-rabbit 594 (Invitrogen Cat #A11010). Nuclei were counterstained with Hoechst 33258 pentahydrate (bis-benzimide) 2 µg/ml (Life Technologies Cat #H3569). Cell death was measured by TUNEL assay (Roche Cat # 11 684 795 910). Images were acquired with fluorescence microscopy utilizing the Leica DMi8. Images were acquired with 10x and 20x objectives. Fields reported in figures are representative of all examined fields.

### Western Blotting

Total protein extracts were prepared from cultures using lysis buffer (50 mM Tris HCl, 100 mM NaCl, 1 mM EDTA, 1% Triton X-100, pH = 7.5) containing protease and phosphatase inhibitor cocktails (Roche). Cell debris was removed by centrifugation, and protein concentration was determined using a standard BSA curve (Pierce, Thermo Scientific Cat #23225). Total protein extracts (30 μg) were loaded onto a NuPAGE 4-12% Bis-Tris gel and electrophoresis was performed in MOPS SDS running buffer (Invitrogen). Proteins were transferred to a PVDF membrane (Invitrogen) and blocked with 5% milk (RPI, Cat #M17200) in PBST (PBS with 0.1% Triton X-100). Incubation with primary antibodies was performed overnight at 4°C. The antibodies used were: p21/CIP1/CDKN1A (R&D systems Cat #AF1074) and ACTB (Cell Signaling Technologies Cat #4967), and HRP-conjugated secondary antibodies (Cell Signaling). Membranes were visualized with enhanced chemiluminescence with SuperSignal West Femto Maximum Sensitivity Substrate (Thermo Scientific Cat #34095) and images were acquired on the ChemiDoc MP Imaging System.

### Statistical analysis

Data are presented as mean ± SEM. All statistical tests were performed using GraphPad Prism 10. The p-value < 0.05 was considered significant difference, determined by Student’s t-test unless otherwise specified. Quantification of fluorescence microscopy were performed using FIJI (ImageJ). Investigators were not blinded to group allocation or outcome assessment. No samples were excluded from this study.

### Single-cell transcriptome data

Single-cell transcriptome data in human colon were retrieved from Wang et al. (2020) using the UCSC Cell Browser tool (Perez et al., 2025).

