## Supplemental Information for "Novel role of the lncRNA EPR as oncosuppressor in intestinal cancer"

### These Authors contributed equally

This document contains:

1) Supplemental figures with legends

2) Supplemental references

#### SUPPLEMENTAL FIGURES

```
Score:214 bits(237), Expect:1e-59,
Identities:385/551(70%),  Gaps:40/551(7%), Strand: Plus/Plus

human  1    TTCCTTCTTGATCCTTCACAAGGGAA---GATTTTCTTTT--TTA-----------GAG-  43
            |||||| | |||||||||||||| ||   | ||||  |||  |||           |||
mouse  43   TTCCTTGTCGATCCTTCACAAGGAAATGAGTTTTTAATTTATTTATTCTTTGCCTTGAGA  102

human  44   GTACAGATTCCTCTTAGTCAAGTCCTGATTAAAACTCCAGCTAAGACATTAGTAAGCCTT  103
            ||||||||||  |||     ||||||| || |||  || || ||||  ||||| ||| |
mouse  103  GTACAGATTCACCTTGAGGGAGTCCTGCTTCAAAG-CCGGC-AAGA--TTAGTGAGCTTA  158

human  104  GGTTAGTGAAGTGGCATCAGGAAGTGCCTACATTTTCATGGCCTGGTAGCGTTCAGT-GA  162
            ||   ||||| | ||| |  |||||  |  ||||| || | ||||| || |  ||   ||
mouse  159  GGACGGTGAACTAGCACCGTGAAGT--CAGCATTTGCAGGACCTGGAAGAGAGCAAAAGA  216

human  163  AAATGTTCATTAACAGACACAGGCCATTCAGTCCCGAATCCCAAGACACTGAAGACTCTG  222
             |||| | | |||| |      || |   ||||| ||  ||||| ||    |||||| |
mouse  217  GAATGCTTACTAACTGC-----GCAACCTAGTCCTGA--CCCAACACGAGAAAGACTGTC  269

human  223  TTTGAATCAGACTCACGGGTTCCTTCCTAGCCACTCTCAGGGACAGGAATGCTTCTGGTG  282
            |   || ||| |||||||| ||| |||| ||  ||||| |||| ||||||| ||||||||
mouse  270  T---AACCAGCCTCACGGGCTCCGTCCTGGCTTCTCTC-GGGAGAGGAATGTTTCTGGTG  325

human  283  AAGAAGTTTTCGGTGGTGGTTCATGGAGCTTCCCTACACCAACTTGGAAATGGCATTCAT  342
            || |      || | ||  | ||||||||| || | ||||||||||||  |||| |||||
mouse  326  AAAACAC---CGTTAGTCTTCCATGGAGCTACCATTCACCAACTTGGAGGTGGCGTTCAT  382

human  343  TTTATTGGCTTTTGTTATCTTTTCCTTATTTACCCTGGCTTCCATCTACACTACTCCGGA  402
            |||| |||||||  | ||||||||||| |||| ||||||||||||||||| |  ||  |
mouse  383  TTTACTGGCTTTCTTCATCTTTTCCTTGTTTACTCTGGCTTCCATCTACACCAGCCCTAA  442

human  403  TGACAGTAATgaagaggaagaacatgaaaaaaagggaagggaaaagaaaaggaaaaagtc  462
            ||| || |||||||| || ||     |   ||||| || |  || |||||| ||| |||
mouse  443  TGAGAGGAATGAAGACGATGACTTCCACCTAAAGGAAAAGAGAAGGAAAAGAAAAGAGTT  502

human  463  tgaaaagaagaaaaaTTGCTCAGAGGAAGAGCACAGAATTGAAGCTGTTGAGCTATGATC  522
            | ||  |||||||||||||||||| ||||| |||| ||||||| |  |  | | |||| |
mouse  503  TAAAGGGAAGAAAAATTGCTCAGATGAAGAACACAAAATTGAAACAATGCAACCATGACC  562

human  523  TCATAGCCACC  533
            ||  |||||||
mouse  563  TC--AGCCACC  571
```

EPR sequence alignment

*Supplementary Figure S1 – Related to Figure 1. Sequence conservation between human EPR and mouse Epr*

Alignment of the 5' region of human and mouse *EPR* sequences using Standard nucleotide BLAST

A

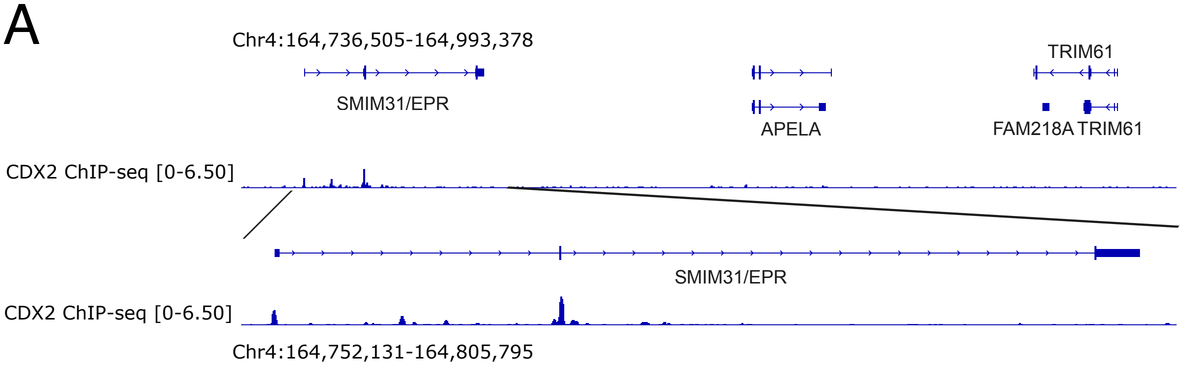


B

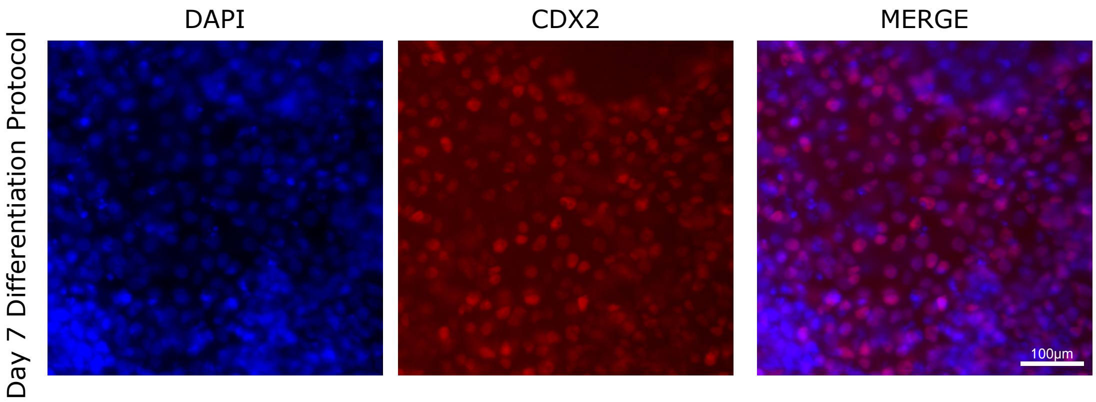


*Supplementary Figure S2 – Related to Figure 2. CDX2 binds the promoter and regulates expression of EPR*

(A) IGV screenshot of the *SMIM31/EPR* locus. Top Panel: Tracks from top to bottom: refseq gene track; CDX2 ChIP-seq in LS180 cells – Data from (Meyer et al. 2012). Bottom Panel: zoom in to the *SMIM31/EPR* gene body.

(B) Representative immunofluorescence staining of CDX2 expression at Day 7 of the differentiation protocol.

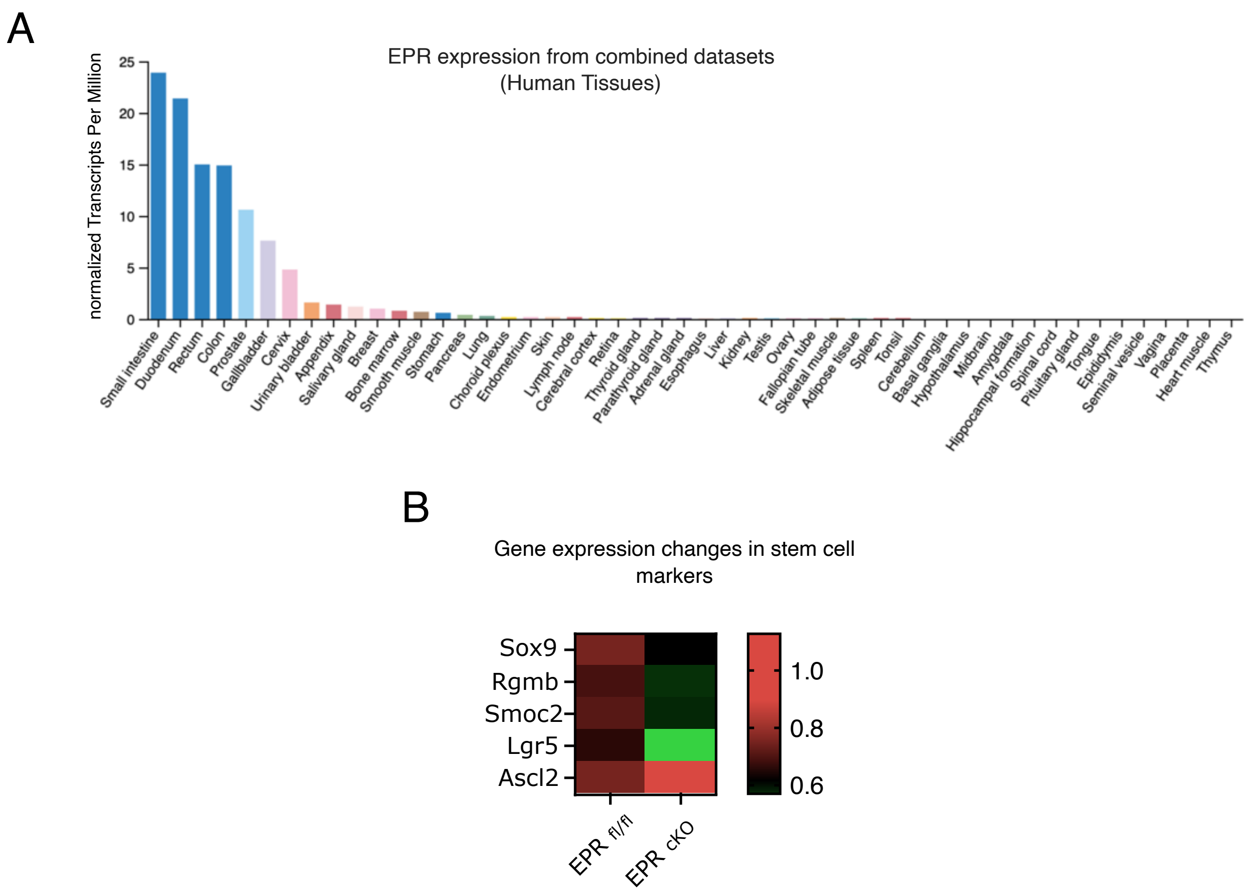


*Supplementary Figure S3 – Related to Figure 3. EPR expression in human*

(A) Expression of human *EPR* (presented as TPM) in different tissues according to The Human Protein Atlas.

(B) Heat map illustrating changes in the expression of genes encoding stem cells markers in *Epr*$^{cKO}$ versus *Epr*$^{fl/fl}$ RNA sequencing.

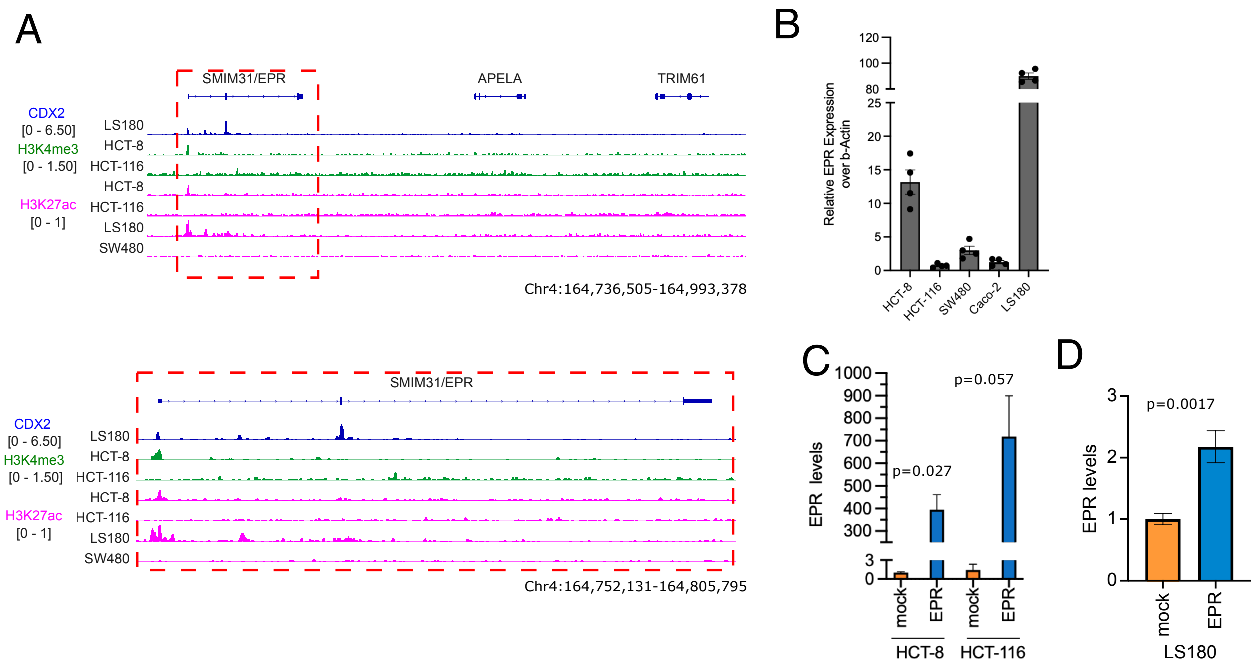


*Supplementary Figure S4 – Related to Figure 5. EPR is expressed in a subset of human colon cancer cell lines*

(A) IGV screenshot of the *SMIM31/EPR* locus. Top Panel: Tracks from top to bottom: refseq gene track; CDX2 ChIP-seq in LS180 cells – data from (Meyer et al. 2012) – blue; H3K4me3 ChIP-seq in HCT-8 and HCT-116 cells - data from (Abraham et al. 2017; Brinkman et al. 2012) – green; H3K27ac ChIP-seq in HCT-8, HCT-116, LS180, and SW480 cells - data from (Abraham et al. 2017; Brinkman et al. 2012; McCleland et al. 2016) – pink. Bottom Panel: zoom in to the *SMIM31/EPR* gene body.

(B) Relative expression of endogenous *EPR* in a panel of colon cancer cell lines.

(C) Relative expression of *EPR* upon transfection of plasmid expressing human *EPR.*
